# Human neurons lacking amyloid precursor protein exhibit cholesterol-associated developmental and presynaptic deficits

**DOI:** 10.1101/2022.12.28.522116

**Authors:** Haylee Mesa, Elaine Y. Zhang, Yingcai Wang, Qi Zhang

## Abstract

Amyloid precursor protein (APP) produces aggregable β-amyloid peptides and its mutations are associated with familial Alzheimer’s disease, which makes it one of the most studied proteins. However, APP’s role in the human brain remains unclear despite years of investigation. One problem is that most studies on APP have been carried out in cell lines or model organisms, which are physiologically different from human neurons in the brain. Recently, human induced neurons (hiNs) derived from induced pluripotent stem cells provide a practical platform for studying the human brain *in vitro*. Here, we generated APP-null iPSCs using CRISPR/Cas9 genome editing technology and differentiate them to matured human neurons with functional synapses using a two-step procedure. During hiN differentiation and maturation, APP-null cells exhibited less neurite growth and reduced synaptogenesis in serum-free but not serum-containing media. We have found that cholesterol (Chol) remedies those developmental defects in APP-null cells, consistent with Chol’s role in neurodevelopment and synaptogenesis. Phenotypic rescue was also achieved by co-culturing those cells with wildtype mouse astrocytes, suggesting that APP’s developmental role is likely astrocytic. Next, we examined matured hiNs using patch-clamp recording and detected reduced synaptic transmission in APP-null cells. This change was largely due to decreased synaptic vesicle (SV) release and retrieval, which was confirmed by live-cell imaging using two SV-specific fluorescent reporters. Adding Chol shortly before stimulation mitigated the SV deficits in APP-null iNs, indicating that APP facilitates presynaptic membrane Chol turnover during SV exo-/endocytosis cycle. Taken together, our study in hiNs supports the notion that APP contributes to neurodevelopment, synaptogenesis and neurotransmission via maintaining brain Chol homeostasis. Given the vital role of Chol in the central nervous system, the functional connection between APP and Chol bears important implication in the pathogenesis of Alzheimer’s disease.

## Introduction

Amyloid precursor protein (APP) is a type I membrane protein with single transmembrane domain which connects a long N-terminal extracellular fragment and a short C-terminal intracellular fragment (Müller et al., 2017). It has drawn tremendous attention because its proteolytic product (β-amyloid peptides, Aβs) makes amyloid plaques, a pathological hallmark for Alzheimer’s disease (AD) (Selkoe and Hardy, 2016). Studies since early 1980s had elucidated how Aβs is carved from APP by β- and γ-secretases (β/γS) sequentially (de Strooper et al., 2010; Selkoe, 1998). However, the majority of APP undergoes nonamyloidogenic cleavage by α- and γ-secretases (α/γS) without producing Aβs. So far, genetic studies about the patients of rare and inheritable forms of AD with early onset (i.e., familial AD, fAD) have identified hundreds of mutations in APP and Presenilin 1/2 (i.e., catalytic subunits of γS), supporting the notion that APP proteolysis and/or its proteolytic products (namely Aβs) are causal factors for fAD (Selkoe and Hardy, 2016). Aβ, soluble or aggregated, are believed to be disruptive to neuronal functionality and survival (Palop and Mucke, 2010).

A powerful tool to study protein function is to remove it from its natural environment, i.e., gene knockout. In early 1990s, three strains of APP-deficient mice were generated (Li et al., 1996; Müller et al., 1994; Zheng et al., 1995). Although viable and fertile, mice completely lacking APP are 15-20% smaller and exhibit various neurological defects including reduced locomotor activity, reactive gliosis, and agenesis of corpus callosum (Li et al., 1996; Müller et al., 1994; Zheng et al., 1995), which suggests developmental and physiological roles for APP. Nevertheless, the underlying mechanism remains unsettled, especially in the aging brain (Zheng and Koo, 2011). For example, study on somatosensory cortex (layers III&V) of adult APP-null mice found an increase in dendritic spine density (Bittner et al., 2009) whereas examining cortical layers II/III and hippocampal CA1 pyramidal neurons of aging APP-null mice found a significant decrease instead (Lee et al., 2010). In addition to the variation between preparations, difference between organisms matters more, especially for complex neurological disorders like AD (Kokjohn and Roher, 2009). Recent development of induced pluripotent stem cells (iPSCs) circumvents ethical concern and leads to a variety of *in vitro* human cell preparations like 2D cultures and 3D organoids for studying aging as well as aging-related disorders (Mertens et al., 2018) like AD (Arber et al., 2017; Israel et al., 2012). One approach is to generate iPSCs from AD patients bearing disease-related genetic mutations or polymorphisms (Amponsah et al., 2021). Alternatively, one can modify AD-associated genes like APP and Presenilin in iPSCs originated from healthy subjects (Essayan-Perez et al., 2019). Fong *et al* showed that human astrocytes differentiated from APP-null iPSCs had reduced lipoprotein uptake, less cholesterol (Chol), and increased expression of genes for Chol synthesis and uptake (Fong et al., 2018). On the other hand, Kwart *et al* found that human neurons derived from APP-knockout iPSCs had significantly smaller endosomes and likely cellular defects in endocytosis pathways (Kwart et al., 2019). However, it remains unclear how the deletion of APP and consequent disruption of Chol homeostasis and endocytosis can impair synaptic function and endanger neural health. Notably, AD is believed to start from synaptic dysfunction that proceeds and correlates to neurodegeneration and cognitive decline (Selkoe, 2002).

The most common form of AD is sporadic or late-onset (sAD or LOAD) (Blennow et al., 2006). Genome-wide association studies have been conducted in large cohorts across populations, nations, continents, and races. Various genetic risk factors but not APP or Presenilins have been identified (Bellenguez et al., 2020; Tanzi, 2012). Those genes are involved in Chol metabolism, cell membrane trafficking, and immune response (Karch and Goate, 2015). Notably, Chol is a great influencer for the latter two (Lingwood and Simons, 2010; Platt et al., 2016) and it has several functional ties to APP. On one hand, APP distribution and cleavage are affected by Chol in the surface and intracellular membranes (Vetrivel and Thinakaran, 2010). On the other hand, APP and its proteolytic fragments modulate Chol uptake and synthesis (Fong et al., 2018; Kwart et al., 2019). More importantly, Barrett et al identified a Chol-interacting motif in APP (Barrett et al., 2012), structurally linking APP and Chol together. However, how APP and Chol collaboratively contribute to AD pathogenesis remains unclear. Here, we use CRISPR/Cas9 genomic editing technology to selectively eliminate APP in human iPSCs. Using a two-step protocol, we differentiate those APP-null iPSCs and its isogenic control to mature and synaptically connected neurons. Using morphological, optical, electrophysiological and pharmacological tools, we examine the developmental and physiological changes in human APP-null neurons. Moreover, we investigate whether and how Chol mediates those phenotypic changes caused by APP knockout.

## Results

### Generation of APP-null iPSCs

To obtain APP-null human neurons (hiNs) form iPSCs, we started with a wild type iPSC line (WT thereafter) that was provided by the NIH Regenerative Medicine Program as wildtype control. Within APP’s exon 3, we selected a twenty-nucleotide sequence for designing sgRNA (**Figure 1a** & **S1**). Synthesized 20-base pair DNA sequence corresponding to this sgRNA was cloned into a CRISPR/Cas9 vector, pSpCas9(BB)-2A-Puro, which co-expresses Cas9 and Puromycin-resistant gene using a viral T2A sequence. Transfected cells were selected against Puromycin and separated to single-cell colonies. Clone 31 was selected using polymerase chain reaction (PCR) with two pairs of primers (i.e., p02/p04 for control and p05/p06 for CRISPR/Cas9 deletion (**Figure 1b** & **S1**)). DNA sequencing of Clone 31 revealed four-nucleotide deletion within APP’s exon 3 (**Figure 1c**) which causes a reading frame shift and hence a premature termination of translation. Using Western blot, we confirmed the absence of APP in clone 31 (**Figure 1d**). To ensure that the genomic editing is limited to APP, we checked the other two members of APP gene family, amyloid precursor-like protein 1 and 2 (APLP1&2), which highly resemble APP in gene sequence and partially redundant in functionality. Western blot showed no significant difference at protein level for both APLP1 and 2 between KO and wild-type iPSCs (**Figure 1d**). Therefore, we conclude that APP is selectively knocked out in the iPSCs clone 31 (KO thereafter).

**Figure 1.**
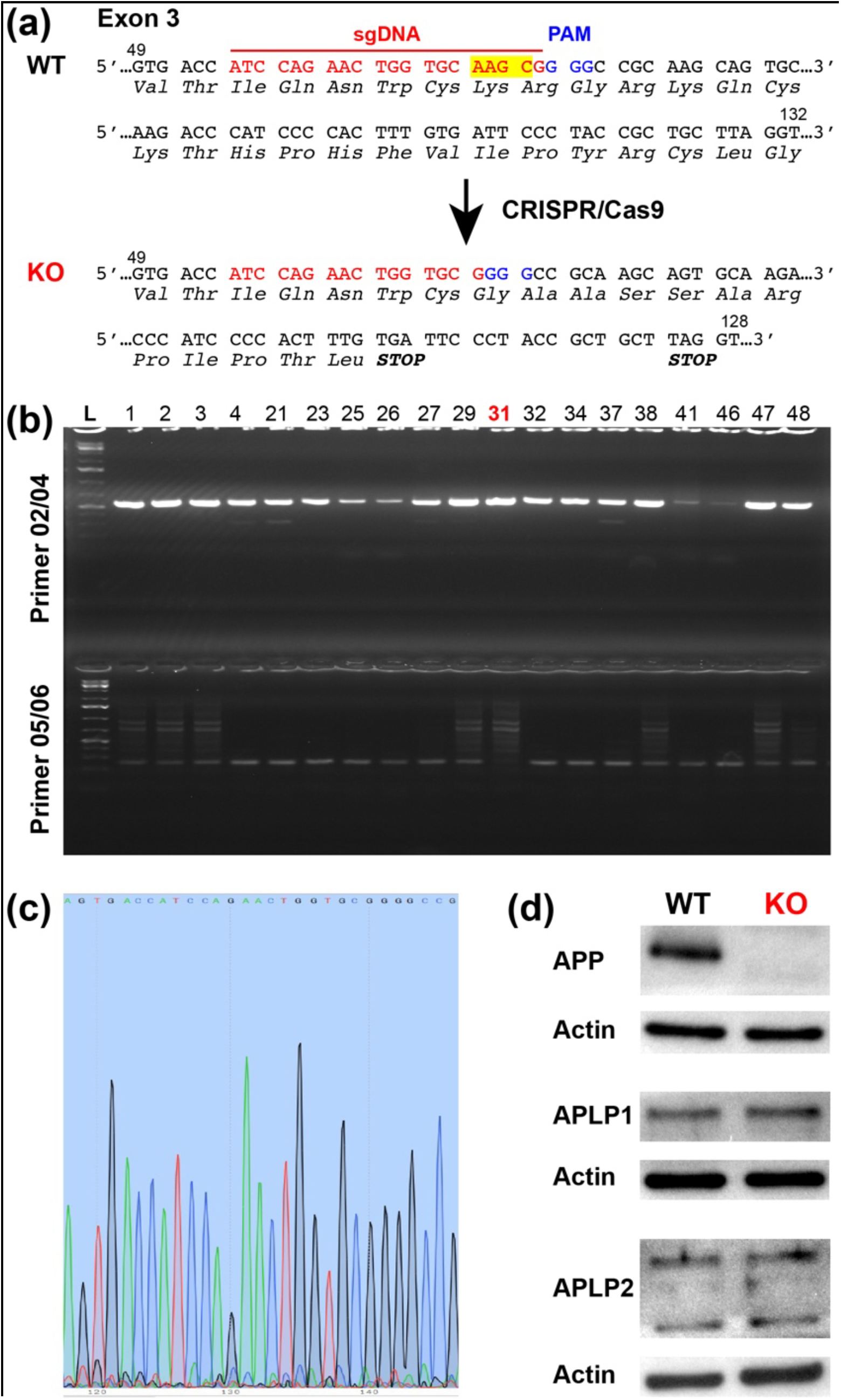
Generation of APP-null human iPSCs. (**a**) sgDNA and targeting sequences (red) are within APP’s exon 3. The nucleotide number is in reference to the beginning of exon 3. Deletion of 4 bp by CRISPR/Cas9 introduces two STOP codons, prematurely terminated APP translation. (**b**) Single clones of resulting iPSCs are assessed by PCR using two pairs of primers P02/P04 and P05/P06. Clone #31 has amplicons with 02/04 but not 05/06, indicating the absence of correct APP transcripts. (**c**) Sequencing of genomic DNA extracted from clone #31 confirmed 4-bp deletion. (**d**) Western blots of APP, APLP1 and APLP2 along with Actin from WT and KO (#31) iPSCs demonstrate the absence of APP but not APLP1 and 2.

### Neurodevelopment of APP-null cells

Since APP reportedly plays a neurotrophic role during development (Müller et al., 2017), we examined the differentiation and maturation of hiNs from KO iPSCs. Based on our previous study (Gu et al., 2015), we utilized a two-step differentiation protocol, i.e., from iPSCs to neuronal precursor cells (NPCs) and then to hiNs (**Figure 2a**). First, we applied neuron induction medium (NIM) to differentiate iPSCs to NPCs. We assessed the differentiation by checking the expression of nestin, an NPC-specific marker (**Figure 2b**). The fraction of nestin-positive cells substantially increased after the application of NIM and peaked between day 4 and 8 (**Figure 2c**), suggesting the completion of the first phase. Notably, there was no significant difference in nestin-positive fractions between KO and WT cells (**Figure 2c**), indicating that APP has little effect on NPC differentiation. Next, we replated both WT and KO NPCs on Matrigel-coated coverslips and fed them with serum-free neurobasal plus medium every three days. We assessed the differentiation of hiNs by monitoring the appearance of Tuj1 (a neuron-specific marker) and glial fibrillary acidic protein (GFAP, an astrocyte-specific marker) (**Figure 2b**). Most NPCs differentiated to Tuj1- or GFAP-positive cells, whose fractions stabilized between day 20 to 30 (**Figure 2d**), suggesting a completion of differentiation. In order to achieve full synaptic connectivity and to simulate the aging brain, we maintained hiNs for at least 90 days. We examined synaptogenesis by reverse transcription quantitative polymerase chain reaction (RT-qPCR) and fluorescent immunostaining for a synapse-specific protein, Synaptophysin (i.e., Syp). RT-qPCR showed that Syp expression rose slowly and plateaued around day 50 and that there was no significant difference between WT and KO cells (**Figure S2**). We also measured the extension and complexity of Tuj1-positive processes (likely neurites) at day 50 using Sholl analysis (Gensel et al., 2010) and found a moderate but significant decrease in APP-knockout cells (**Figure 2e&f**), in good agreement with APP’s moderate contribution to neurodevelopment. Moreover, we found a significant difference between WT and KO when we measured the density of Syp (i.e., the number of Syp puncta per 20 μm Tuj1-positive processes) at day 50. It was significantly lower in KO cells (**Figure 2g**), suggesting that the lack of APP caused a reduction in synaptogenesis.

**Figure 2.**
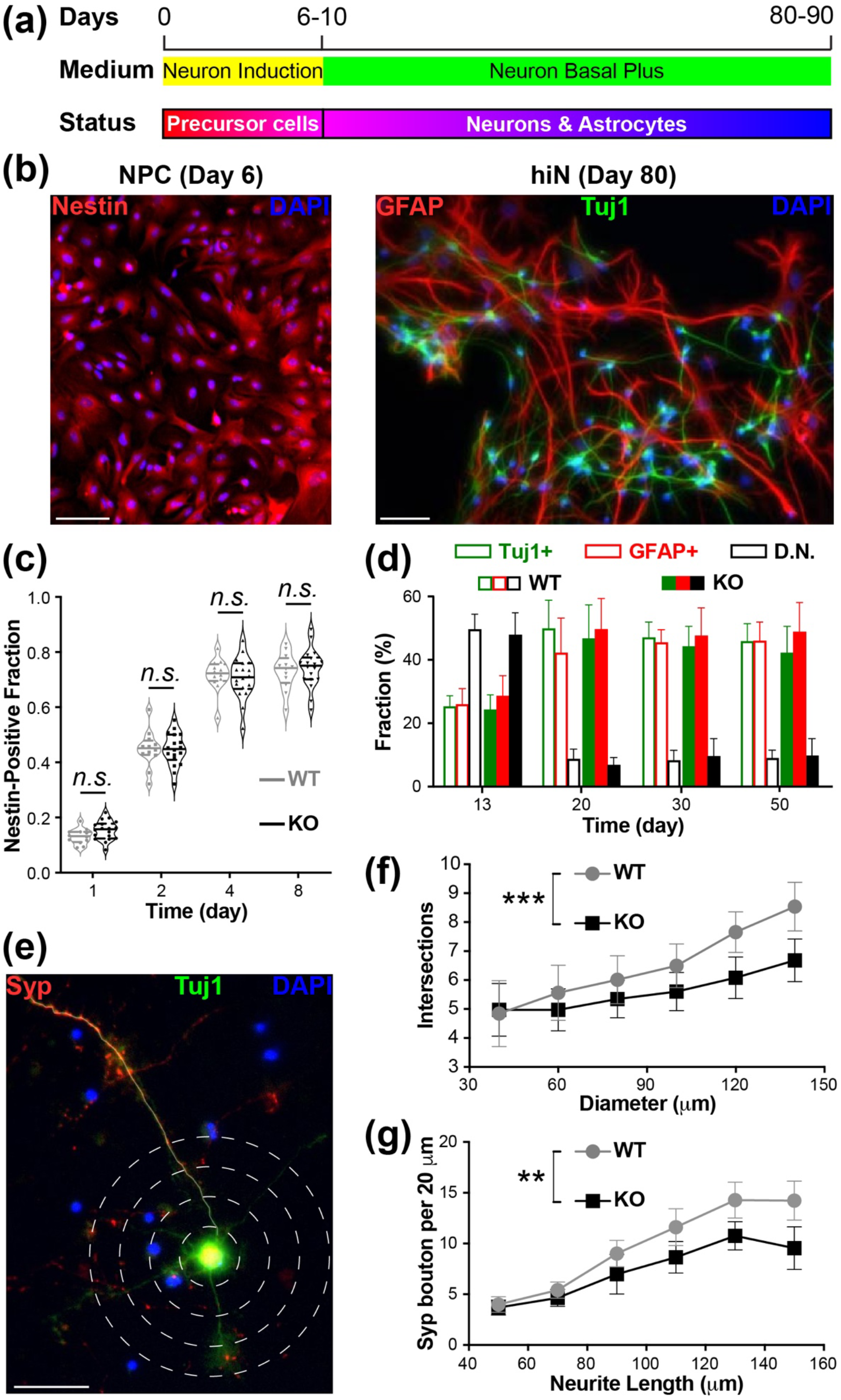
Differentiation of human iPSCs to hiNs and detection of neurodevelopmental defects in KO cells. (**a**) Two-step differentiation procedure. (**b**) Sample images of immunofluorescence labeling of WT cells for Nestin (red) at the end of the first step (left); and labeling for GFAP (red) and Tuj1 (green) at the end of the second step (right). Nuclear staining by DAPI (blue) is used to count all cells. Scale bar, 100 μm. (**c**) Violin plot with medians (solid lines) and quartiles (dash lines) shows a progressive appearance of Nestin-positive cells until they plateaued by day 4-8 in the first step. There are no significant differences (*n.s.*) between WT (gray) and KO (black) cells at all four dates. All n = 15 FOVs (i.e., 5 FOVs per batch and 3 batches per group) for both WT and KO at four dates. Two-tailed Student’s *t*-test for comparing WT and KO at every date, all *p* > 0.05. (**d**) Bar plot for the cell fractions of Tuj1+ (green), GFAP+ (red), and double negative (D.N.) cells at four dates during the second step. There are no significant differences (*n.s.*) between WT (open) and KO (filled) groups for all three staining types at all dates. All n = 15 FOVs (i.e., 5 FOVs per batch and 3 batches per group) for both WT and KO at four dates. Two-tailed Student’s *t*-test, all *p* > 0.05. (**e**) Sample image of immunofluorescence labeling of WT cells (at 50 days into the second step) for Tuj1 (green, to illustrate neurites) and Synaptophysin (Syp, red), with DAPI marking all cell nuclei. The concentric dash-line circles with a 20-μm interval are used for Sholl analysis. The semi-transparent solid white line illustrates one of the neurites (by Tuj1 staining). Scale bar, 50 μm. (**f**) Mean and error plot of intersection counts between neurites and concentric circles at 50 days into the second step. There is a significant difference in the total intersection counts between WT and KO cells. Both n = 18 hiNs (i.e., 6 hiNs per batch and 3 batches per group) for WT and KO. Two-tailed Student’s *t*-test, *p* < 0.001 (***). (**g**) Mean and error plot of Syp bouton counts (per 20 μm along neurites) in the same samples as (**f**). There is a significant difference in the average counts at all lengths between WT and KO cells. Both sample sizes are the same as those in (**f**). Two-tailed Student’s *t*-test, *p* < 0.01 (**).

To improve neuronal differentiation and synaptogenesis in the second step, we tested serum-containing media widely used for primary neuronal culture (Liu and Tsien, 1995). Although adding serum made no difference for the resulting fractions of neurons (Tuj1+) and astrocytes (GFAP+) (**Figure 3a**), we did find that neurite growth and synaptogenesis in KO cells became similar to those of WT control at day 50 (**Figure 3b** & **c**). Alternatively, we experimented purified wild-type murine astrocytes as feeder cells based on our previous work with embryonic stem cells (Gu et al., 2015). We plated iPSC-derived NPCs on mouse cortical astrocytes and observed similar improvements in KO cells (**Figure 3a – c**). It is clear that both serum and wildtype astrocytes brought the neurodevelopment of KO cells to the same level as that of WT. The most likely candidate is Chol in the form of high-density lipoprotein (HDL) in serum or from astrocytes. Chol is known to be crucial for axon growth and synaptogenesis (Pfrieger, 2003a). To test that, we first applied 10 μg/mL HDL on to KO cells right after plating the iPSC-derived NPCs. We measured neurites growth and Syp puncta density at day 50. There was a significant improvement by both counts (**Figure 3d & e**). Next, we used 0.5 mM Chol-saturated methyl-β-cyclodextrin (Chol:MβCD), often used to supplement Chol to the plasma membrane (Zidovetzki and Levitan, 2007). Again, it significantly increased neurite growth and synaptogenesis (**Figure 3d & e**). Together, our data suggest that the moderate developmental deficits in KO cells are mediated by Chol, although we cannot rule out other factors that is redundant or complimentary to Chol. Since serum-containing media has to be used with mitosis inhibitor to avoid glia overgrowth (Liu and Tsien, 1995), we used serum-free media and wildtype murine astrocytes as feeder cells for the rest of this study.

**Figure 3.**
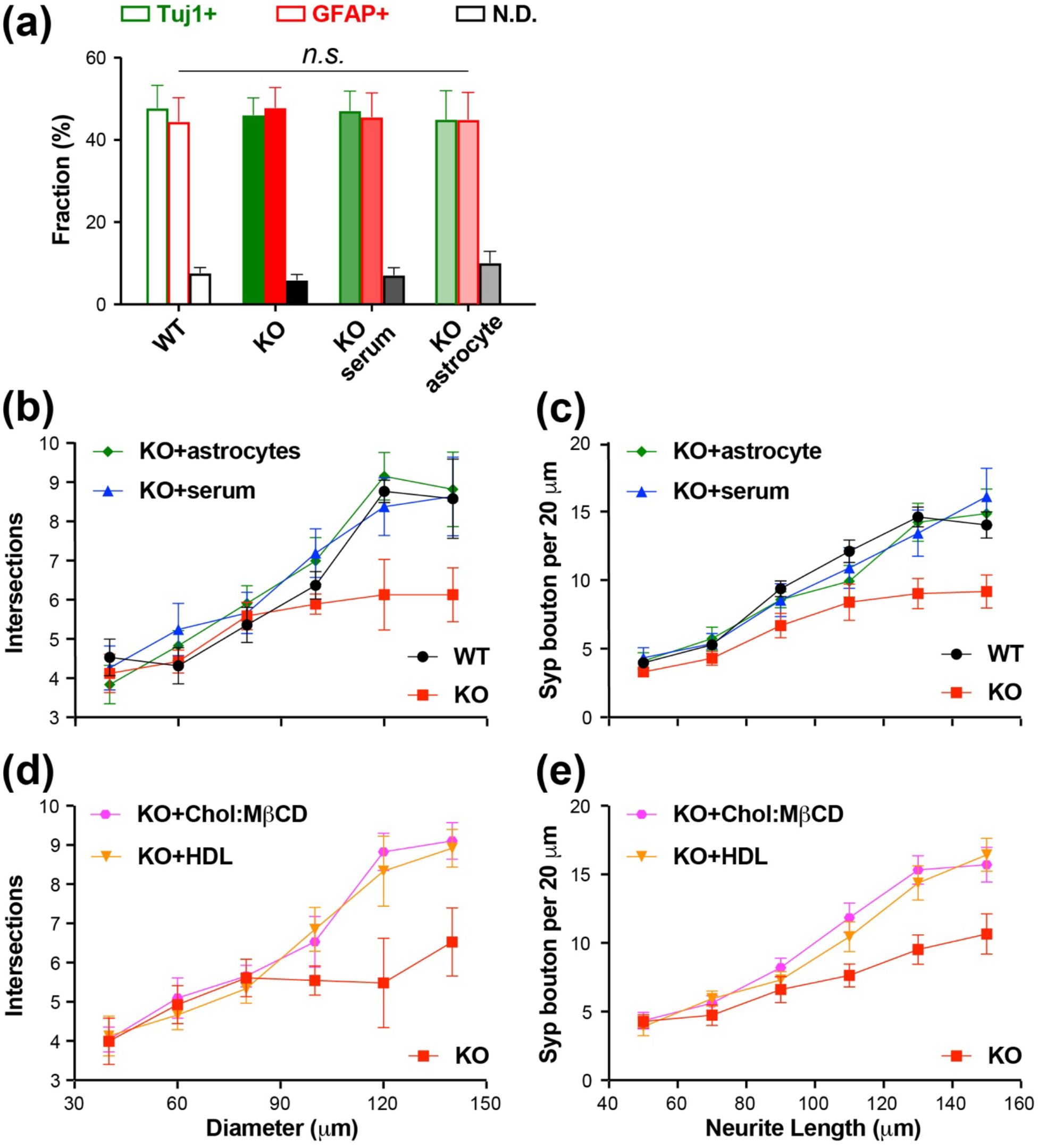
Chol rescues KO’s neurodevelopmental defects. (**a**) Bar plot shows that adding serum or astrocytes during the second step makes no significant difference (*n.s*.) on the fractions of differentiated cell-types. All n = 15 FOVs (i.e., 5 FOVs per batch and 3 batches per group) for the four groups. ANOVA and post-hoc Tukey-Kramer test between WT and each KO groups, all *p* > 0.05. (**b**) Mean and error plot shows that adding serum or astrocytes to KO group rescues neurite growth deficits. All n = 18 cells (i.e., 6 cells per batch and 3 batches per group) for the four groups. Comparison are done using the summations of all data points for every condition. Two-tailed Student’s *t*-test, *p* < 0.01 for KO *vs*. WT but both *p* > 0.05 for KO + serum or KO + astrocytes *vs*. WT. (**c**) Mean and error plot shows that adding serum or astrocytes to KO group improves synaptogenesis. All sample sizes are the same as those in (**b**). Two-tailed Student’s *t*-test, *p* < 0.005 for KO *vs*. WT and *p* > 0.05 for KO + serum or KO + astrocytes *vs*. WT. (**d**) Mean and error plot shows that adding HDL or Chol:MβCD to KO group improves neurite growth. All n = 18 cells (i.e., 6 cells per batch and 3 batches per group) for the four groups. By two-tailed Student’s *t*-test, both *p* < 0.005 in comparison to KO. (**e**) Mean and error plot shows that adding HDL or Chol:MβCD to KO group increases synaptogenesis. All sample sizes are the same as those in (**d**). Two-tailed Student’s *t*-test, both *p* < 0.001 in comparison to KO.

### Presynaptic changes in APP-null neurons

To study the electrophysiological character of KO cells, we performed whole-cell voltage-clamp recordings after 90 days in differentiation. 17 out of 19 randomly selected cells in the KO group possessed a voltage-dependent and tetrodotoxin (TTX)-sensitive inward current (**Figure 4a**), suggesting the presence of voltage-gated Na^+^-channels (VGSCs). This is comparable to the WT group (12/13 cells recorded). Since neurotransmitter release is one of the hallmarks for functional neurons, we measured the activity-evoked and Ca^2+^-dependent release of synaptic vesicles (SVs) in both groups using FM1-43, a green styryl dye commonly used to label SVs (Betz et al., 1992). For loading, hiNs were exposed to Tyrode’s solution containing 90 mM K^+^ and 10 μM FM1-43 for 2 minutes, and then washed with dye-free normal Tyrode’s solution (4 mM K^+^) for 5 minutes. FM1-43 puncta were evident along the processes of hiNs (**Figure 4b1**). When challenged with 10-Hz, 2-min electric field stimulation, there were stimulation-induced FM1-43 fluorescence loss in both (**Figure 4b2**), which are similar to that of cultured mouse hippocampal neurons. We also recorded spontaneous excitatory postsynaptic currents (sEPSCs) in both WT and KO groups (**Figure 4c**), demonstrating that those cells were synaptically connected. Together with previous immunocytochemistry results, we conclude that both groups of cells were indeed functional neurons with functional synapses.

**Figure 4.**
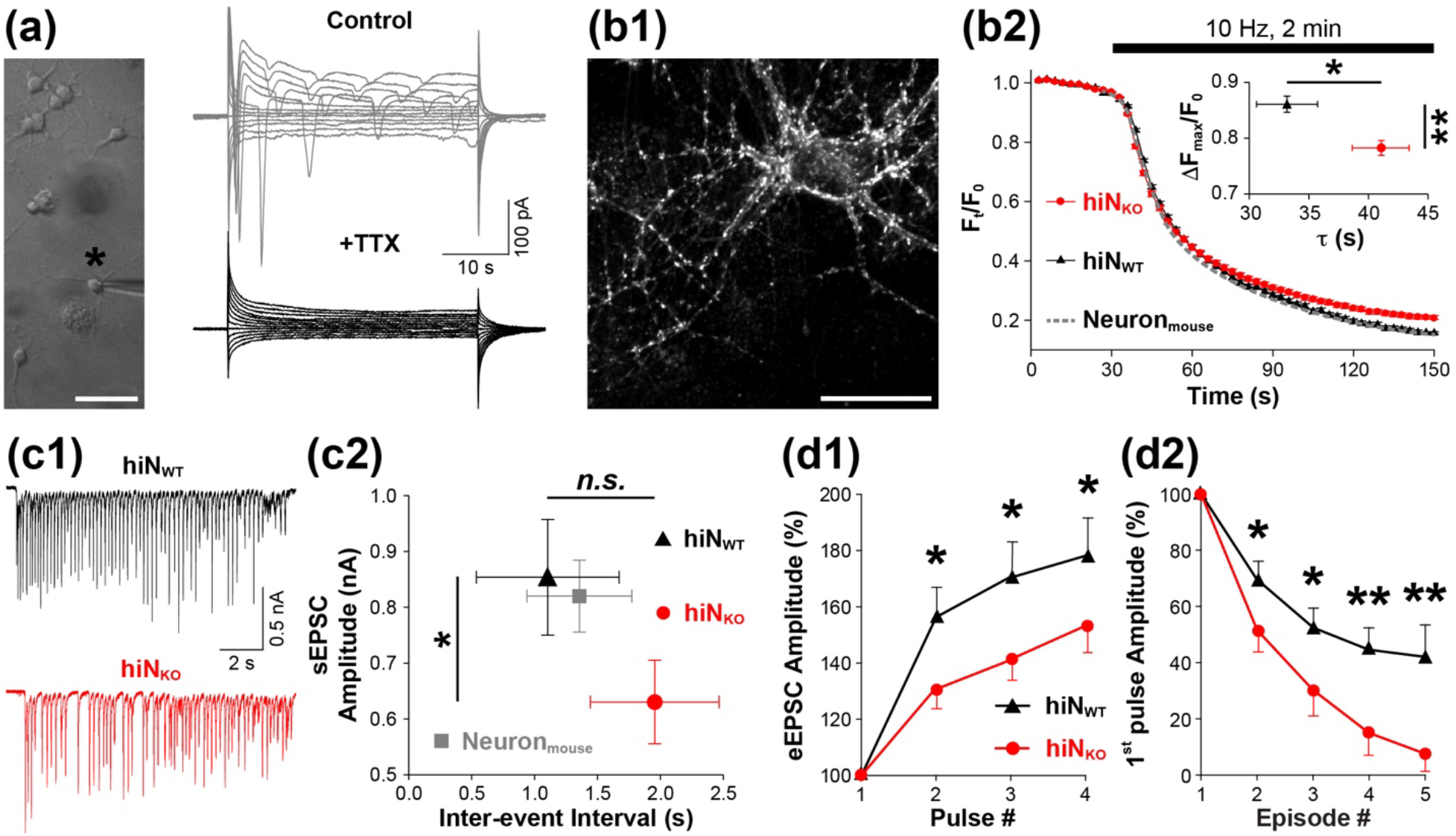
Mature KO hiNs exhibits reduced excitatory neurotransmission. (**a**) Sample image (left) and records (right) of whole-cell patch clamp recording in a WT hiN. Scale bar, 100 μm. Voltage-dependent inward current is detected (upper right), and it is blocked by TTX (lower right), suggesting the presence of voltage-gated sodium channels in hiNs. (**b1**) Sample image of hiNs loaded with FM1-43 whose fluorescent puncta indicates presynaptic terminals. Scale bar, 20 μm. (**b2**) Mean and error plot shows KO hiNs (hiN_KO_, red circles) have slower and less FM1-43 destaining than WT hiNs (hiN_WT_, black triangles) or wild-type mouse hippocampal neurons (Neuron_mouse_, dotted line) during a 10-Hz, 2-min electric field stimulation (indicated by black bar). Inset is a 2D mean and error plot for average maximal FM1-43 florescence decrease (ΔF_max_/F_0_) and average decay time constant (τ) between KO and WT hiNs, showing significant less florescence decrease and slower decay in KO hiNs. Both n = 180 FM1-43 puncta (i.e., 20 randomly selected FM1-43 puncta per FOV, 3 FOVs per batch, and 3 batches per group). Two-tailed Student’s *t*-test, *p* < 0.01 (**) for ΔF_max_/Foand *p* < 0.05 for τ(*). (**c1**) Sample recordings of sEPSCs in WT (black) and KO (red) hiNs. (**c2**) 2D mean and error plot shows sEPSCs in KO hiNs (red circle) exhibit decreased amplitude but similar frequency (i.e., longer inter-event interval) in comparison to WT hiNs (hiN_WT_, black triangle). There is a significant difference in sPESC amplitudes between WT and KO but not in inter-event interval. Data from wild-type mouse hippocampal neurons is used as reference. All n = 450 events (i.e., 30 consecutive events per cell, 5 cells per batch, and 3 batches per group). Two-tailed Student’s *t*-test for KO *vs*. WT, *p* < 0.05 (*) for the amplitudes and *p* > 0.05 (*n.s*.) for the inter-event interval. (**d1**) Mean and error plot shows the average of four pluses in all five stimulation episodes between KO (red circle) and WT (black triangle) hiNs. The amplitude increases of the 2^nd^, 3^rd^, and 4^th^ pluses are significantly less in KO. Both n = 18 hiNs (i.e., 6 hiNs per batch and 3 batches per group) for WT and KO. Two-tailed Student’s *t*-test, all three *p* < 0.05 (*). (**d2**) Mean and error plot shows the average of the first pluses across all five stimulation episodes between KO and WT hiNs. The amplitude decreases of the first pluses in episodes 2, 3, 4 and 5 are significantly more in KO. Both sample sizes are the same as those in (**d1**). Two-tailed Student’s *t*-test, *p* < 0.05 (*) for the first pulses in 2^nd^ and 3^rd^ episodes, and *p* < 0.01 (**) for the first pulses in 4^th^ and 5^th^ episodes.

Notably, KO hiNs show moderate but significant reduction as well as slow down of FM1-43 destaining, along with smaller sEPSC amplitude than the WT (**Figure 4b & c**). To understand such change, we measured the evoked EPSCs (eEPSCs) by applying five episodes of 0.5-Hz 8-sec (i.e., 4 pulses) field stimulation with a 10-second interval between consecutive episodes. We used 10-second interval because it was long enough for fast vesicle recycling but not enough for conventional clathrin-mediated endocytosis (Alabi and Tsien, 2013; Wu et al., 2014). Within the 4-pulse stimuli, we observed short-term facilitation of eEPSC amplitudes in both WT and KO groups. However, there was significantly less facilitation in the KO (**Figure 4d1**). When comparing the first eEPSCs across the five episodes, we found a depression in both groups but significantly more in KO hiNs (**Figure 4d2**). These results indicate that KO cells exhibit defective synaptic transmission and short-term plasticity, most likely at presynaptic terminals.

Such short-term eEPSCs facilitation (within every episode) and depression (across consecutive episodes) could result from changes in intracellular Ca^2+^ concentration ([Ca^2+^]_i_), SV pool size, and SV release probability. First, we checked evoked Ca^2+^-influx using a low-affinity and cellpermeant Ca^2+^-indicator, Fluo-4·AM, ideal for measuring fast [Ca^2+^]_i_ entry (Gee et al., 2000; Grienberger and Konnerth, 2012). To identify synaptic boutons, we retrospectively loaded cells with FM4-64, a red styryl dye selectively labeling SVs (Rouze and Schwartz, 1998; Vida and Emr, 1995). In order to eliminate spontaneous neuronal firing and evoked postsynaptic Ca^2+^-influx, we applied NBQX and D-AP5 to block AMPA and NMDA receptors respectively. We continuously imaged Fluo-4 fluorescence before, during and after a 5-s 10-Hz electric field stimulation (**Figure 5a1**). We did not detect any difference in the peak values or half rise times (t_1/2_) of Fluo-4 fluorescence at synaptic boutons upon stimulation (**Figure 5a2**). After stimulation, the decay constant (τ) of the Fluo-4 fluorescence was the same (**Figure 5a2**). Those results suggest that APP knockout did not affect presynaptic Ca^2+^-influx or post-stimulation [Ca^2+^]_i_ recovery. Can APP deletion change presynaptic resting [Ca^2+^]_i_? To test that, we used Fura-2·AM, a high-affinity ratiometric Ca^2+^ indicator suitable for measuring low [Ca^2+^]_i_ (Grynkiewicz et al., 1985; Miyawaki et al., 1997). Again, FM4-64 staining was used to mark synaptic boutons. We applied ionomycin to permeabilize hiNs and equalized their [Ca^2+^]_i_ using Tyrode’s solutions with defined Ca^2+^concentrations. As such, we generated a working curve for Fura-2 ratio (R_340/380_) *vs*. [Ca^2+^]_i_ at synaptic boutons. We found no significant difference in the presynaptic resting [Ca^2+^]_i_ between WT and KO cells (67.5 ±4.8 nM and 70.9 ±4.3 nM respectively) (**Figure 5b**).

**Figure 5.**
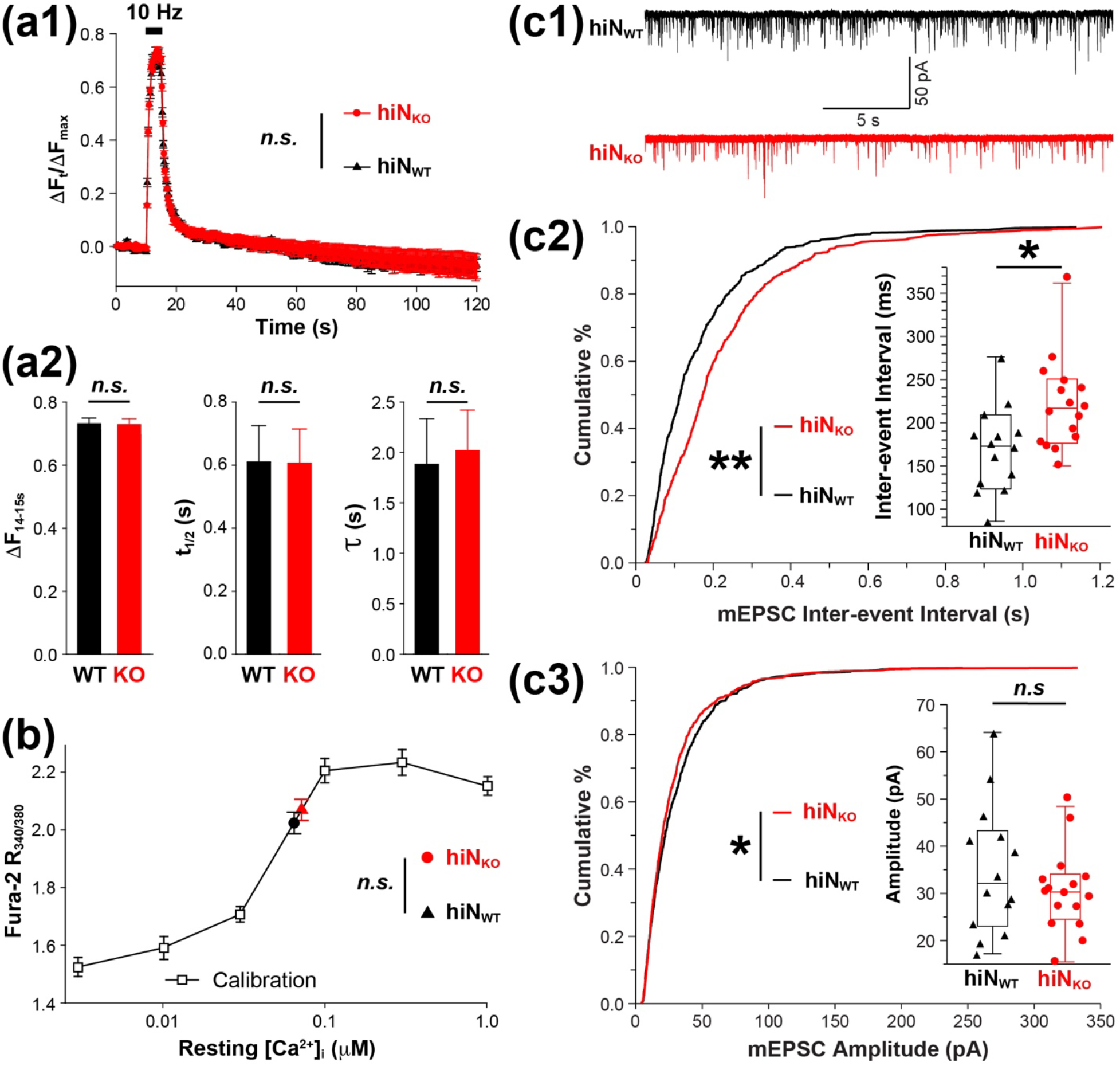
APP knockout affects neurotransmitter release but not intracellular Ca^2+^ at presynaptic terminals. (**a1**) Mean and error plot of presynaptic (determined by FM4-64 staining) Ca^2+^ imaging, i.e., relative fluorescence change of Fluo-4 (ΔF_t_/ΔF_max_), before, during, and after 5-s 10-Hz electrical stimulation (11^th^ - 15^th^ seconds, black bar). In term of total fluorescence increase during the stimulation, no significant difference is detected between KO (red circle) and WT (black triangle). Both n = 180 FM1-43 puncta (i.e., 20 randomly selected FM1-43 puncta per FOV, 3 FOVs per batch, and 3 batches per group). Two-tailed Student’s *t*-test, *p* > 0.05 (*n.s*.). (**a2**) Bar plots show no significant differences in peak response (at 14^th^ - 15^th^ seconds), in response speed (i.e., half rising time during stimulation, t_1/2_), and in recovery speed (i.e., fluorescence decay time constant after stimulation, τ) are detected between KO (red) and WT (black). Both n = 180 FM1-43 puncta (i.e., 20 randomly selected FM1-43 puncta per FOV, 3 FOVs per batch, and 3 batches per group). Two-tailed Student’s *t*-test, all *p* > 0.05 (*n.s.*). (**b**) Mean and error plot of ratiometric Ca^2+^ imaging using Fura-2 (R_340/380_) shows intracellular Ca^2+^ concentration ([Ca^2+^]_i_) calibration curve (black open square) and presynaptic (marked by FM4-64 staining) [Ca^2+^]_i_. The presynaptic resting [Ca^2+^]_i_ of WT and KO are estimated to be 66.8 ± 5.1 nM and 71.1 ±3.9 nM respectively. No significant difference is detected between KO and WT. For calibration, n = 90 hiNs (i.e., 10 randomly selected hiNs per FOV, 3 FOVs per batch, and 3 batches in total). For KO and WT, both n = 180 FM1-43 puncta (i.e., 20 randomly selected FM1-43 puncta per FOV, 3 FOVs per batch, and 3 batches per group). Two-tailed Student’s *t*-test, *p* > 0.05 (*n.s.*). (**c1**) Sample mEPSC recordings of WT (black) and KO (red). Cumulative distribution plot of inter-event interval (i.e., inverse of frequency) (**c2**) and cumulative distributions of mEPSC amplitude (**c3**) show significant difference in mEPSC frequency but not amplitudes between WT (black) and KO (red). Both n = 750 events (i.e., 50 consecutive events per cell, 5 cells per batch, and 3 batches per group) for KO and WT. *Kolmogorov-Smirnov* test, *p* < 0.01 for inter-event interval and *p* < 0.05 for amplitude. Insets in (**c2**) and (**c3**) are box plots with all data points (i.e., average values of individual cells) of KO (red circles) and WT (black triangles). According to the average values of individual cells, there is a significant difference in frequencies but not amplitudes between KO and WT.

To further evaluate synaptic change in KO cells, we measured miniature EPSCs (mEPSCs), whose amplitudes and frequencies are often used to evaluate post- and presynaptic changes respectively (Zucker and Regehr, 2002). Between WT and KO cells, the most significant difference was the mEPSC frequency (i.e., inter-event interval, sample records shown in **Figure 5c1**). We detected significantly longer inter-event intervals in the KO when counting mEPSC events (50 consecutive events from every recording) or the average of average frequency for every recorded cell (**Figure 5c2** and insert respectively). As lower mEPSC frequency (i.e. longer inter-event interval) generally indicates reduced Pr,v. This result indicates presynaptic deficiency in SV release for KO cells. In case of mEPSC amplitudes, there was no significant difference between the two groups (**Figure 5c3** and inset showing average of averages), arguing against a significant postsynaptic change in the KO group.

### APP KO has less releasable synaptic vesicles and slower recycling

To directly examine the effect of APP KO on presynaptic terminals especially SVs, we used Synaptophysin-pHTomato (SypHTm) to label SVs. Synaptophysin is SV-specific and pHTomato is a red fluorescence protein sensitive to pH. The latter is inserted in the luminal domain of the former, so that pHTm can report deacidification and reacidification following SV exo-/endocytosis respectively (Li and Tsien, 2012). Notably, pHTm weakly fluoresces at low pH, which makes synaptic boutons visible even at rest. To mobilize all releasable SVs, we again applied 10-Hz 2-min electric stimulation; then, we sequentially applied 50mM NH_4_Cl that neutralized all SVs and pH5.5 Tyrode’s solution that quench cell surface SypHTm (**Figure 6a**). Expectedly, we detected (1) a rapid increase of SypHTm fluorescence representing evoked SV exocytosis, (2) a subsequent decrease representing compensatory endocytosis and reacidification, (3) maximal SypHTm fluorescence by 50mM NH_4_Cl, and (4) minimal SypHTm fluorescence by pH5.5 Tyrode’s solution. We normalized the fluorescence changes at individual boutons to the maximal (as 1.0) and the minimal (as 0.0) which were set by NH_4_Cl and pH5.5 treatments respectively. Hence, the fractions of SypHTm at the presynaptic surface (*i*), in the releasable pool (*ii*) and in the total pool (*iii*) (**Figure 6a**) are all measurable. Among the three fractions, only (*ii*) was significantly lower (~15 % less than WT) in the KO hiNs (**Figure 6b**), suggesting a reduction of the releasable pool. Although there was no significant difference in the rapid increase of pHTm fluorescence, there was a difference in the decay phase (i.e., τ_WT_ = 41.8 ± 2.3 s, τ_KO_ = 49.2 ±3.7 s; **Figure 6c**). This result suggests a slower retrieval and/or recycling of released SVs.

**Figure 6.**
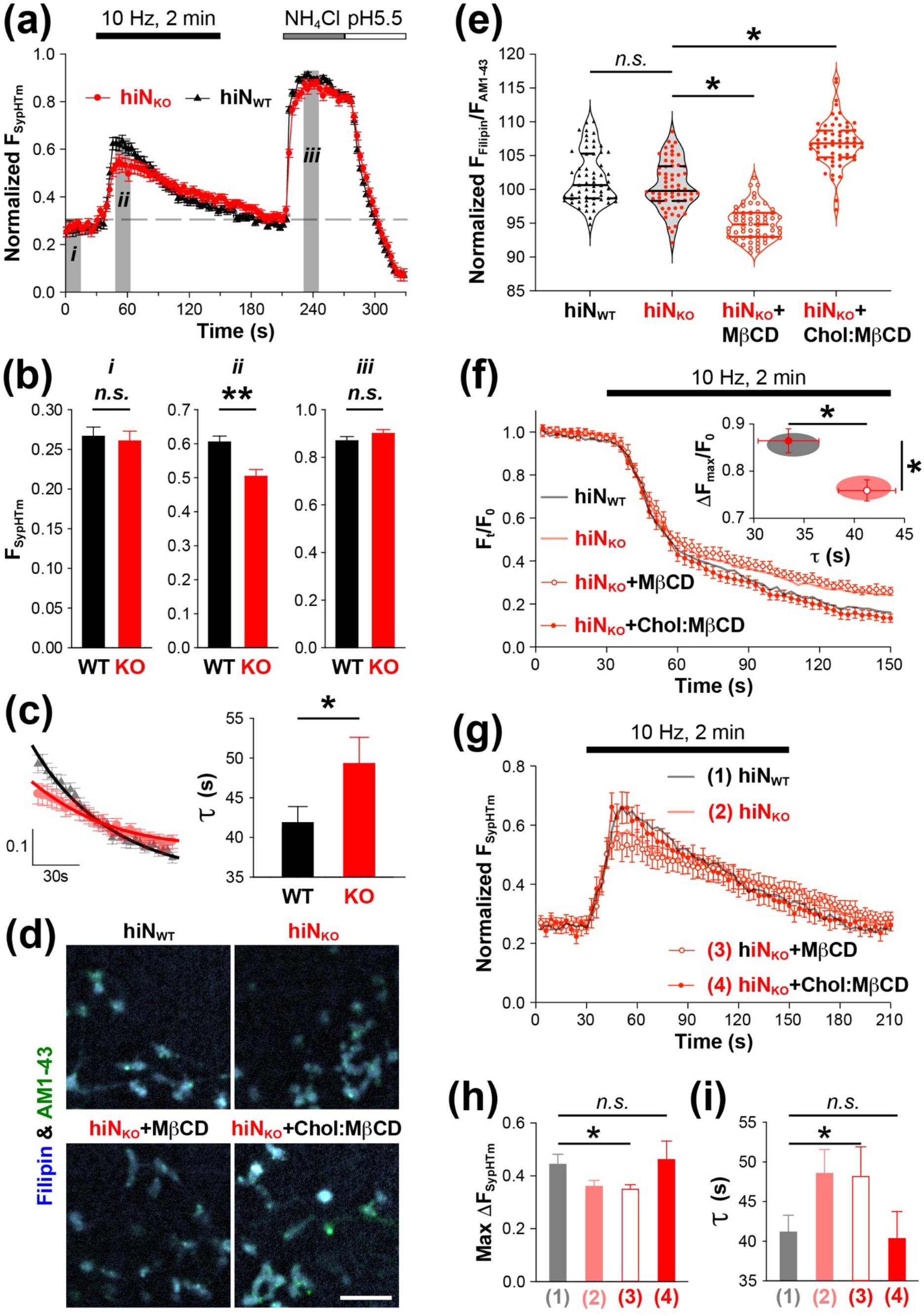
APP knockout reduces SV turnover via disrupting Chol. (**a**) Mean and error plot shows SypHTm fluorescence intensity change during 10-Hz 2-min electrical field stimulation, which is normalized to maximal (by 50 mM NH_4_Cl) and minimal (by pH5.5 Tyrode’s solution) fluorescence. KO (red circles) showed less increase and slower recovery than WT (black triangles). Both n = 180 SypHTm puncta (i.e., 20 randomly selected puncta per FOV, 3 FOVs per batch, and 3 batches per group) for KO and WT. (**b**) Bar plots of average values from three periods (i.e., shaded areas, i, ii, and iii, in **a**) demonstrates that KO (red) exhibits significantly less SypHTm fluorescence responses than WT (black) only during stimulation (period ii). Same sample sizes as (**a**). Two-tailed Student’s *t*-test, *p* > 0.05 (*n.s*.) for i and iii and *p* < 0.01 (**) for ii. (**c**) The left is single-exponential decay fittings for average SypHTm fluorescence changes between 61^th^ and 150^th^ second of the imaging (i.e., during the last 90 seconds of the stimulation). KO (red) is slower than WT (black). The right is bar plot of the average decay time constants (τ) shows that KO is significantly longer than WT, suggesting slower SV recycling. Same sample sizes as (**a**). Twotailed Student’s *t*-test, *p* < 0.05 (*). (**d**) Sample images of Filipin (blue) and AM1-43 (red) costaining in WT and KO hiNs as well as KO hiNs treated with MβCD and Chol: MβCD. Scale bar, 10 μm. (**e**) Violin plot with medians (solid lines) and quartiles (dash lines) shows synaptic (based on AM1-43 puncta) membrane Chol estimated by Filipin and AM1-43 fluorescence ratio (normalized to the average ratio in WT hiNs as 100%). There is no significant difference (*n.s*.) between WT (black) and KO (red) hiNs. In comparison to KO hiNs, there is significant decrease or increase in KO hiNs treated with MβCD or Chol: MβCD respectively. All n = 60 ROIs (i.e., 10 ROIs per FOV, 3 FOVs per batch and 2 batches per group) for all four groups. Two-tailed Student’s *t*-test for comparing WT and KO (*p* > 0.05) and for comparing KO and KO with MβCD or Chol: MβCD treatments (both *p* < 0.05). (**f**) Mean and error plot shows FM1-43 destaining by 10-Hz 2-min electrical field stimulation, which is normalized to pre-stimulation baseline (F_0_). KO acutely treated with MβCD (open red circles) showed little difference from untreated KO cells (semi-transparent solid red line) whereas adding Chol: MβCD (filled red circles) improves KO cells to be like WT (semi-transparent solid black line). n = 180 FM1-43 puncta (i.e., 20 randomly selected puncta per FOV, 3 FOVs per batch, and 3 batches per group) for every group. Inset is mean and error plot of average relative FM1-43 detaining and time constant from individual FOVs (i.e., n = 9). Same symbols for MβCD and Chol: MβCD treated KO hiNs; semi-transparent red and black ovals represent the ranges (i.e., mean ±SEM) of KO and WT respectively. Two-tailed Student’s *t*-test, *p* < 0.05 (*) for ΔF_max_/F_0_ and τ between MβCD and Chol: MβCD treated KO hiNs. (**g**) Mean and error plot shows SypHTm fluorescence changes by 10-Hz 2-min electrical field stimulation, which is normalized in the same way as (**a**). Again, acute MβCD application (open red circles) had little effect in comparison to KO alone (semi-transparent solid red line) whereas adding Chol: MβCD (filled red circles) makes KO cells behave like WT (semi-transparent solid black line). n = 180 FM1-43 puncta (i.e., 20 randomly selected puncta per FOV, 3 FOVs per batch, and 3 batches per group) for every group. (**h**) Bar plot of maximal SypHTm increase during the stimulation shows that Chol: MβCD but not MβCD recovers the response back to WT level. Same n as (**f**). ANOVA and post-hoc Tukey-Kramer test between WT (1) and KO with MβCD (3) or with Chol: MβCD (4), *p* < 0.05 (*) for (1) *vs*. (3) and *p* > 0.05 (*n.s*.) for (1) vs. (4). (**i**) Bar plot of SypHTm fluorescence recovery time constant shows that Chol: MβCD but not MβCD recovers the time constant back to WT level. Same n as (**f**). ANOVA and post-hoc Tukey-Kramer test between WT (1) and KO with MβCD (3) or with Chol: MβCD (4),*p* < 0.05 (*) for (1) *vs*. (3) and *p* > 0.05 (*n.s*.) for (1) vs. (4).

Since APP directly interacts with Chol (Barrett et al., 2012), since Chol is important for exo- and endocytosis (Linetti et al., 2010; Yue and Xu, 2015), and since co-cultured wild type astrocytes amended chronic Chol deficits, we hypothesize that the presynaptic changes in KO hiNs is also related to Chol. But it is likely about acute Chol change instead of Chronic alternation because Filipin staining of synaptic membrane Chol showed no significant difference between WT and KO hiNs (**Figure 6d&e**). To test if APP deletion is related to acute membrane Chol change at synapses, we applied MβCD or Chol:MβCD 10 minutes before stimulation to quickly decrease or increase membrane Chol. Notably, we used relatively high concentrations of MβCD and Chol:MβCD (3 and 2 mM respectively) to compensate for the short application duration. Filipin staining for membrane Chol showed that such acute MβCD or Chol:MβCD treatment caused synaptic membrane Chol reduction or increment respectively (**Figure 6d&e**). Using FM1-43 imaging, we found that Chol:MβCD brought the FM1-43 destaining kinetic in KO hiNs back to the WT level whereas MβCD had little effect if not causing further reduction (**Figure 6f**). Similarly, SypHTm imaging also showed that Chol:MβCD but not MβCD amend the difference between KO and WT hiNs (**Figure 6g-i**). Together, these results suggested that the absence of APP unbalanced the membrane Chol and consequently impairs SV release and recycle. Since the plasma membrane at axon terminal is the most likely membrane section affected by the acute application of Chol:MβCD, we can further conclude that the reduction of Chol in the presynaptic surface membrane in KO cells is the major cause of reduced SV turnover.

## DISCUSSION

APP is one of the most studied protein due to its proteolytic product (i.e., Aβs) and its close tie to AD. While the genetics, biochemistry, and cleavage of APP are well understood, the intrinsic functionality of APP, especially in the adult brain, is far from clear (Zheng and Koo, 2006). Studies of APP-knockout mice revealed its moderate impact on neurodevelopment and synaptic transmission, but the underlying mechanisms remain elusive (Müller and Zheng, 2012). To better understand APP’s role in the human brain and its implication in AD pathogenesis, recent knockout studies utilizing human iPSCs and CRISPR/Cas9-genome editing suggest APP’s relevance to Chol regulation in astrocytes (Fong et al., 2018) and endosome size in neurons (Kwart et al., 2019). However, it remains unanswered if and how APP’s relationship with Chol and endocytosis is relevant to neurodevelopment and synaptic function, especially in the context of AD and neurodegeneration in general. Using genome editing, we produced human iPSCs lacking APP expression. Using a refined two-step procedure, we differentiated APP-knockout iPSCs and the isogenic controls to NPCs and then to matured and synaptically connected hiNs. During the differentiation and maturation, we observed moderate but significant neurodevelopmental deficiency and presynaptic defects. By manipulating Chol, we have demonstrated that those chronic and acute phenotypes can be mostly mitigated by increasing Chol supply and cell membrane Chol content respectively. Based on our morphological and physiological data, and based on the fact that Chol is vital to neurons and synapses, we conclude that Chol dysregulation is at least one of the explanations for the phenotypes of APP-null cells. Furthermore, we propose that disruption of neuronal Chol homeostasis, especially at axon terminals, causes synaptic dysfunction and endangers neuronal survival. In other words, neuronal Chol is likely a converging point of various AD risk factors associated with Chol metabolism and endocytosis.

At the molecular level, different mechanisms can mediate APP’s influence on neuronal Chol. APP isoforms containing the KPI domain at its N-terminal can bind to lipoprotein receptor-related protein 1 (Knauer et al., 1996; Kounnas et al., 1995), which is responsible for neuronal Chol uptake. The cytoplasmic part of APP is linked to multiple LDL receptors by an adaptor protein, FE65 (Pietrzik et al., 2004; Trommsdorff et al., 1998). Additionally, APP has diverse intracellular and extracellular binding partners including SORLA (Schmidt et al., 2007) and ApoE (Haβ et al., 1998), again indicating its potential to influence Chol uptake. More importantly, APP possesses a Chol interactive motif, which is within the Aβ sequence and partially overlaps with APP’s transmembrane domain (Barrett et al., 2012). This structural feature enables APP to interact with membrane Chol (Beel et al., 2010; DelBove et al., 2019; Song et al., 2014, 2013). In our previous study, we showed that point mutations disrupting APP-Chol interaction chronically altered the distribution of synaptic membrane Chol (DelBove et al., 2019). The results from this study further demonstrate that APP also affects synaptic membrane Chol in an acute manner. Therefore, APP can chronically and actually modulate neuronal Chol in developing as well as mature neurons.

While possessing about a quarter of total body Chol, brain metabolizes and regulates Chol independently from other parts of the body due to the separation by blood-brain brain (Dietschy and Turley, 2004). Chol is crucial for the development, functionality and survival of neurons in the brain (Segatto et al., 2014; Zhang and Liu, 2015). Almost all brain Chol is synthesized in astrocytes and supplied to neurons in the form of HDL (Dietschy and Turley, 2004). Intriguingly, ApoE is the major brain lipoprotein and ApoE4 (the highest genetic risk factor for sAD) less intends to form HDL than E2 and E3, indicating a higher burden for maintaining neuronal Chol homeostasis for ApoE4 carriers (Jeong et al., 2019; Leduc et al., 2010; Mahley, 2016). In neurons and almost all mammalian cells, the plasma membrane possesses the majority of cellular Chol whereas intracellular membrane has much less Chol content (Yeagle, 1985). The maintenance of such membrane Chol gradient involves active membrane trafficking and many membrane proteins (Enrich et al., 2015; Schroeder et al., 1996; Yeagle, 1985). On the other hand, disturbance of membrane Chol can affect a lot of membrane proteins (including ion channels, receptors, and transporters in neurons) and cellular signaling (including neurotransmitter secretion) (Jeremic et al., 2006; Koudinov and Koudinova, 2001; Pfrieger, 2003b; Pfrieger and Ungerer, 2011). Clearly, it is extremely important for neurons to maintain Chol gradient between the surface and intracellular membranes. This is particularly challenging at axons and axon terminals where vesicle trafficking and membrane exchanging are the most active (Bloom and Morgan, 2011). Our results agree well with the idea that APP modulates synaptic membrane Chol homeostasis. Thus, APP deletion or mutation may disrupt neuronal membrane Chol and consequently synaptic function as well as neuronal survival. While guided by the popular amyloid hypothesis, previous studies on fAD mutations or sAD risk factors mostly focused on Aβ production and aggregation. With new findings linking APP to Chol, it is time to ask if and how those genetic changes alter Chol metabolism and homeostatic regulation in neurons and other brain cells. And the advances in iPSC and genome-editing technologies have paved way to tackle these questions in a more clinically relevant way.

## Author Contributions

Q.Z. designed and supervised all experiments. H.M. and E.Y.Z. carried out imaging experiments and analyze the data. H.M. made cell cultures, differentiated hiNs, collected samples, performed tests. Y.W. generated APP knockout iPSCs and validated them. Q.Z. also conducted imaging experiments, electrophysiology recording, and data analysis. Q.Z. wrote the paper with the help and inputs from all authors.

## Acknowledgment

We thank R. Blakely for advice and comments and O. Pelletier for technical support. We thank all members of the Zhang Lab for their input and support.

## Abbreviations

Aβ: β-amyloid peptide
AD: Alzheimer’s disease
α/β/γS: α/β/γ-secretases
fAD: familial Alzheimer’s disease
sAD: sporadic Alzheimer’s disease
APLP: amyloid precursor-like protein
APP: amyloid precursor protein
[Ca^2+^]_i_: intracellular calcium concentration
Chol: cholesterol
Chol:MβCD: Chol-saturated methyl-β-cyclodextrin
D.N.: double negative
eEPSCs: evoked excitatory postsynaptic currents
FOV: field of view
GFAP: glial fibrillary acidic protein
HDL: high-density lipoprotein
hiN: human induced neuron derived from induced pluripotent stem cell
iPSC: induced pluripotent stem cell
KO: knock-out
LOAD: late-onset Alzheimer’s disease
MβCD: methyl-β-cyclodextrin
mEPSCs: miniature excitatory postsynaptic currents
NPC: neuronal precursor cells
RT-qPCR: reverse transcription quantitative polymerase chain reaction
sEPSCs: spontaneous excitatory postsynaptic currents
SV: synaptic vesicle
Syp: Synaptophysin
SypHTm: Synaptophysin-pHTomato
TTX: tetrodotoxin
VGSC: voltage-gated sodium channel
WT: wild-type

## Materials and Methods

### Genomic editing and clone selection

The wild-type human iPSC line ND50022 was obtained from NINDS Human Cell and Data Repository. To disrupt APP gene, we designed sgRNA (5’ – ATCCAGAACT GGUGCAAGCG – 3’) and cloned it into CRISPR/Cas9 vector, pSpCas9(BB)–2A–Puro (62988, Addgene). The resulting plasmid was transfected into the wild-type human iPSCs using Lipofectamine (CMAX00015, ThermoFisher Scientific). Transfected iPSC clones were selected against 5 μg/ml Puromycin (A1113802, ThermoFisher Scientific). Puromycin-resistant iPSCs were dissociated by Accutase (07922, STEMCELL Technologies) and plated in very low density for single clones. After one passage, fractions of viable clones were collected and their genomic DNAs were extracted. To identify APP–null iPSC clones, we used two pairs of PCR primers: p02 (5’ – CACTCCACCT GTACCTTACA GT – 3’) & p04 (5’ – GCATAGCGTA TCTACTAAGA GTAC – 3’) and p05 (5’ – ATCCAGAACT GGTGCAAGCG – 3’) & p06 (5’ – TACTGCTCCT ATAGGGTCAG TGCA – 3’). Their relative positions in the APP cDNA sequences are shown in Figure S1. PCR amplification with both pairs were done with 55°C annealing and 45s extension. Clone(s) positive for p02/p04 but negative for p05/p06 were selected for further characterization.

#### Animals and astrocyte culture

All animal handling and procedures were carried out in accordance with the Care and Use of Laboratory Animals of the National Institutes of Health and approved by the Florida Atlantic University Institutional Animal Care and Use Committee. Pregnant (E14) C57BL/6J (wild-type) female mice were purchased from Jackson Laboratory. After arrival, those mice were acclimated at FAU vivarium until giving birth. Postnatal cortical cultures were prepared as previously described (Liu and Tsien, 1995) Briefly, mouse cortices were collected from postnatal day 0-2 new born pups and dissociated into a single-cell suspension with a 10-min incubation in 0.01% Trypsin-EDTA (Life Technologies) followed by gentle trituration using three glass pipettes of different diameters (~ 1, 0.5, and 0.2 mm), sequentially. Dissociated cells were recovered by centrifugation (x 200 g, 5 minutes) at 4 °C and re-suspended in media composed of Minimal Essential Medium (MEM, Life Technologies) with 10%/vol fetal bovine serum (Omega). 5 mL of cell suspension was plated in T75 culture flasks precoated with Matrigel (Corning). Cells were allowed to settle down for 2 hours before the addition of 10 mL media. After 7-10 days in vitro, cells (mostly astrocytes) reached confluency whereas neurons die out. Cells in the flasks were washed three times with ice-cold PBS and shaken repeatedly during every wash. To ensure the depletion of neurons, oligodendrocytes, and microglia, flasks were filled with cold PBS and placed in a culture shaker (180rpm) for at least 2 hours. Then, cells in the flasks were washed with ice-cold PBS for another three times with stronger tapping in between. The absence of neurons and other glial cells were confirmed by RT-qPCR. Purified astrocytes in the flasks were detached by 0.01% Trypsin-EDTA and collected as cell pellets after (x 200 g, 5 minutes) at 4 °C. After wash and spin down, purified astrocytes were resuspended in serum-free Neurobasal Plus medium (ThermoFisher Scientific) and plated on 12mm-ⵁ round glass coverslips (~200,000 cells/mL) precoated with Matrigel (1 coverslip per well in 24-well plates) for more than 1 hour. After two hours, we added 1mL media for every coverslip and maintained them until the plating of NPCs.

#### iPSCs culture and their differentiation to neurons and astrocytes

Human iPSCs with or without CRISPR/Cas9-based genome editing were grown in 35mm culture dish until ~70% confluency. After three times wash with pre-warmed PBS, cells were detached by Accutase (ThermoFisher Scientific), and replated in new 35mm dishes or 6-well plates precoated with Matrigel (~1,000 cells/mL). For storage, iPSC pellets were resuspended in mFreSR™ Cryopreservation media (STEMCELL Technologies) and placed in −20°C for 2 hours, −80°C for overnight and finally in liquid nitrogen for long-term storage. For the 1^st^ step of iPSC differentiate, Neural Induction Medium (STEMCELL Technologies) was applied in replated iPSCs and exchanged every other day for 6-10 days. In the second step of differentiation, cells were washed and detached as previous described. The detached cells were plated on Matrigel-coated 12mm-ⵁ round glass coverslips (~200,000 cells/mL) or mouse cortical astrocytes monolayers and fed with Neurobasal Plus medium every three days until reach 80-90 days. For HDL application, HDL purchased from VWR (Cat. No. BIRBORB82274-1) was made into 1mg/mL suspension in PBS as stock solution and the HDL stock solution was added to culture media with 1:100 dilution ratio.

#### RT-qPCR

Every step of the procedure after sample preparation, including RNA extraction, cDNA synthesis, qPCR and analysis, was performed blind. Cells were collected at the defined time points and stored in Ambion RNAlater (Thermo Fisher Scientific, Waltham, Massachusetts) at 4 °C for no more than two weeks. RNA was extracted using the Aurum Total RNA minikit (skipping the DNase step) (Bio-Rad, Hercules, California) and immediately processed or stored at 80 °C. QuantiTect Reverse Transcription Kit (Qiagen) was used for genomic DNA removal and cDNA synthesis according to the provided instructions. cDNA was stored at −20 °C. Quantitative PCR was performed using Maxima SYBR Green qPCR Master Mix (Thermo Scientific), 96-well clear plates (Thermo Scientific), and optical flat 8-cap strips (Bio-Rad) on an Opticon 2 from MJ Research with Opticon Monitor 3 software (BioRad). Reactions were set up manually using an 8-channel pipette (30-300μL) for the master mix and a single channel P2 pipette for the cDNA samples. Appropriate controls, including a reverse transcriptase-negative sample, a no template control, and a pooled cDNA sample (taken from cultured hippocampal neurons), were used on every plate to ensure consistency. Information about the primers, including sequences, sequence accession number, and annealing temperature used, can be found in **Table S1**. Optimal annealing temperature under the conditions described was experimentally determined by running a temperature gradient. The program for qPCR is as follows: (1) 95 °C for 15 minutes, (2) 95 °C for 10 seconds, (3) in designated annealing temperature (**Table S1**) for 30 seconds, (4) 72 °C for 30 seconds, (5) plate read, (6) repeat steps 2 - 5 44 times, (7) 72 °C for 10 minutes, (8) melting curve from 50 °C to 95 °C and read every 0.2 °C, (9) 72 °C for 10 minutes, (10) 4°C for storage. C(t) values were exported analyzed using Microsoft Excel and SigmaPlot. The 2^-ΔC(t)^ method (Schmittgen and Livak, 2008) was used to quantify the expression of the target genes relative to the expression of three reference genes, β-actin, CypA, and Ppp1ca, which were selected based on their consistent expression in neurons and astrocytes (Bonefeld et al., 2008). The 2^-C(t)^ values were first normalized to the average 2^-C(t)^ of all samples of the gene, or intra-normalized, before further quantification.

#### Immunocytochemistry, Fluorescence image and Analysis

For immunostaining, coverslips were fixed in PBS containing 4% paraformaldehyde, washed, blocked for 1 h with PBS containing 1% BSA, and incubated overnight at 4 °C with primary antibodies. Secondary antibodies with distinct fluorophores were then incubated at room temperature for 2 hours. Fluorescence imaging was performed on a Nikon Ti-E inverted microscope equipped with 20X Apo and 100X (N.A. = 0.75 and 1.40, respectively) objective and a Flash4.0 sCMOS camera (Hamamatsu). The optical filter sets used are from Chroma or Semrock. For each fluorescence channel, images were taken with the same acquisition settings (i.e., excitation intensity and exposure time). For Filipin staining, cells were loaded with a fixable styryl dye, AM1-43, during the treatments to label cell membranes. After stimulation and imaging, samples were then washed with PBS and fixed by 4% paraformaldehyde (PFA) for 30 minutes at room temperature. PFA were neutralized by 1.5 mg/mL glycine and cells were washed three times with PBS. Next, those fixed cells were incubated with PBS containing 50 μg/mL Filipin and 10% fetal bovine serum at room temperature for 2 hours. Stained cells were washed before mounted on glass slides. Both Filipin fluorescence (blue) and AM1-43 fluorescence (green) images for the same fields of view were acquired. For image analysis, Fluorescence images were thresholded in ImageJ and used as a mask with the same thresholding value applied across different image sets. The masks were subjected to minimum particle size restrictions and ROI sets were generated for every image sets. These ROI sets were applied to the original image and mean intensities were measured. Background subtraction was performed on intensity values measured from every FOV by subtraction of the average of at least 4 different ROIs manually selected from cell free areas. Data were pooled for analysis and total number of ROIs analyzed for every test are stated in the figure legends.

#### Electrophysiology

For every test, we used three different batches of cultures, randomly chose at least five coverslips per batch, and recorded at least one neuron per coverslip. Whole-cell voltage clamp recordings were performed using a Multi-Clamp 700B amplifier, digitized through a Digidata 1440A, and interfaced via pCLAMP 10 software (all from Molecular Devices). All recordings were performed at room temperature. Cells were voltage clamped at −70 mV for all experiments. Patch pipettes were pulled from borosilicate glass capillaries with resistances ranging from 3 - 6 MΩ when filled with pipette solution. The bath solution (Tyrode’s solution) contained (in mM): 150 NaCl, 4 KCl, 2 MgCl_2_, 2 CaCl_2_, 10 N-2 hydroxyethyl piperazine-n-2 ethanesulphonic acid (HEPES), 10 glucose, pH 7.35. The pipette solution contained (in mM): 120 Cesium Methanesulfonate, 8 CsCl, 1 MgCl_2_, 10 HEPES, 0.4 ethylene glycol-bis-(aminoethyl ethane)-N,N,N’,N’-tetraacetic acid (EGTA), 2 MgATP, 0.3 GTP-Tris, 10 phosphocreatine, QX-314 (50 μM), pH 7.2. For the recordings of eEPSCs, a pair of platinum wires were positioned at the edge of the recording chamber and connected to a Grass SD9 stimulator set at 70mV and 1ms duration for single pulses. For the recording of mEPSCs, bath solution was supplied with 1 μM tetrodotoxin (TTX, Abcam). The last 100 seconds of the recording were collected. For event detection of sEPSCs and mEPSCs, we used a template-based method, in which the templates were the average of thousands of sEPSC and mEPSC events randomly selected from hundreds of records collected from different batches of controls. We did not use the threshold-based event detection because the increase of sEPSC or mEPSC amplitudes might lead to more events being larger than an arbitrary threshold and thus an artificial increase of event frequency. The template was generated from our own representative data. All signals were digitized at 20 kHz, filtered at 2 kHz, and analyzed offline with Clampfit software (Molecular Devices).

#### Live Cell Fluorescence Imaging and Image Analysis

Live cell imaging was performed using the same Nikon Ti-E microscope setup previously described. For Ca^2+^-imaging, cells were pre-incubated with Fura-2·AM or Fluo-4·AM (30 minutes, 1 μM, Biotium) at 37°C with 5% CO_2_. After dye loading, coverslips were washed and mounted in an RC-26G imaging chamber (Warner Instruments) bottom-sealed with a 24 × 40 mm size 0 cover glass (Fisher Scientific). The chamber was fixed in a PH-1 platform (Warner Instruments) fixed on the MP-78 stage and bath solutions containing 10 μM NBQX and 20 μM D-AP5 were applied via gravity perfusion with a constant rate of ~50 μL/sec. All perfusion lines were merged into an SHM-6 in-line solution heater (Warner Instruments). The temperatures of both the imaging chamber and the perfusion solution were maintained at 34°C by a temperature controller (TC344B, Warner Instruments). For FM dye loading of the evoked pool of synaptic vesicles, mounted coverslips were incubated with 10 μM FM1-43 or FM4-64 (Life Technologies) for 2 minutes in high K^+^ bath solution containing (in mM): 64 NaCl, 90 KCl, 2 MgCl_2_, 2 CaCl_2_, 10 N-2 hydroxyethyl piperazine-n-2 ethanesulphonic acid (HEPES), 10 glucose, 1 μM TTX, pH 7.35. After FM loading, cells were washed with normal Tyrode’s solution containing 10 μM NBQX and 20 μM D-AP5 for 5 or 10 minutes respectively. Electrical field stimulation (10 Hz, 70V) was triggered by a 5-V 2-ms TTL pulse generated by Clampex software 30 seconds after imaging began, and delivered via a pair of platinum wires attached to both sides of the imaging chamber by a Grass SD9 stimulator. Synchronization of perfusion with image acquisition was via a VC-6 valve system (Warner Instruments) and controlled in Clampex. For SypHTm imaging, the exposure time was 100 ms for all images. For Fluo-4 and FM imaging, the exposure time was 50 ms and the acquisition rate was 1 Hz. For Fura-2, two excitation filters, 340/20 and 380/20 were combined with a Prior L200 light source to excite Fura-2. For calcium concentration calibration, cells were permeabilized with ionomycin and incubated with Tyrode solutions containing designated concentration of calcium. The exposure time was 50 ms with an EM gain of 900 for all images. All images were taken with the same acquisition settings within the same measurements (laser intensity, exposure time, and EM gain).

Image analyses were performed in ImageJ. Four rectangular ROIs were drawn in cell-free regions in every FOV and their intensities averaged. For every type of fluorescence imaging including Fura-2, Fluo-4, FM1-43, and SypHTm, we pooled all background ROIs regardless of transgene differences in order to calculate the mean and standard deviation of the background intensity. Again, a masked threshold approach was applied in ImageJ, and the mean intensity plus two standard deviations was used as the common threshold for all images or image stacks. For every FOV, ROIs were generated by particle analysis based on the binary threshold mask image. FM and SypHTm, watershed segmentation and particle size limits (0.3 – 3 μm) were applied in ImageJ to isolate ROIs representing synaptic boutons (~1 μm). For Fura-2, Fluo-4, and FM time-lapse imaging, the average intensity from 4 background ROIs was subtracted from the average intensity of each individual ROI in the same FOV. Normalization was performed using the average intensity of the first 5 frames.

#### Statistical analysis

All experiments were carried out in a double-blind fashion and repeated with at least three different batches of cultures (N ≥ 3). Every imaging test was repeated in at least three randomly picked coverslips per batch with randomly chosen FOVs per coverslip. No statistical methods were used to predetermine sample size. All values presented are mean ± s.e.m. All fluorescence intensity values are background subtracted. Unpaired two-tailed Student *t*-tests or *Kolmogorov-Smirnov* test were used for two-group comparison. For more than three groups in one measure, statistics were calculated using Two-Way ANOVA with repeated measures followed by the unpaired two-tailed Student *t*-tests. *Kolmogorov-Smirnov* tests were used to compare data of cumulative distributions.

