## Supplemental Table and Figures for "Human neurons lacking amyloid precursor protein exhibit cholesterol-associated developmental and presynaptic deficits"

**Title:**

**Running Title:**

APP regulates Chol for neurodevelopment and synapses

**Authors:**

Haylee Mesa[1†], Elaine Y. Zhang[1,2†], Yingcai Wang[3], Qi Zhang[1,4*]

**Supplementary content:**

Supplementary Table S1

Supplementary Figures and Legends

### 1. Supplementary Table

#### Table S1. qPCR primers

| Primer name | Primer sequence (5' to 3') * | Amplicon Length | $T_M$ (°C) | Annealing T (°C) | mRNA refseq | Gene ID |
|---|---|---|---|---|---|---|
| **Synaptophysin FW** | TGCAGAACAAGTACCGAGAG | 297 bp | 60.4 | 57 | NM_012664.1 | 24804 |
| **Synaptophysin RV** | CTGTCTCCTTGAACACGAACC | | 62.6 | 57 | | |
| **CypA FW** | TATCTGCACTGCCAAGACTGAGTG | 126 bp | 64.6 | 60 | NM_017101.1 | 25518 |
| **CypA RV** | CTTCTTGCTGGTCTTGCCATTCC | | 64.6 | 60 | | |
| **β Actin FW** | AGGCCCCTCTGAACCCTAAG | 118 bp | 64.5 | 60 | NM_031144.3 | 81822 |
| **β Actin RV** | CCAGAGGCATACAGGGACAAC | | 64.5 | 60 | | |
| **Ppp1ca FW** | ATGAGTGTGCCAGCATCAAC | 100 bp | 60.4 | 60 | NM_031527 | 24668 |
| **Ppp1ca RV** | CAGTTGAAGCAGTCGGTGAA | | 60.4 | 60 | | |

*Sources for the primers:

Synaptophysin (Teresa Hsi, ATRC Reagent Bank, Harvard, modified for rat synaptophysin), CypA primers (Langnaese et al., 2008), β-actin (Langnaese et al., 2008), Ppp1ca (Wheeler et al., 2012).

### 2. Supplementary Figures

```
                                            p02
agcacttctggtcccaagcattttggataagggacactccacctgtaccttacagtggaggcttgttagatgctt base pairs
tcgtgaagaccagggttcgtaaaacctattccctgtgaggtggacatggaatgtcacctccgaacaatctacgaa 226 to 300
                                                         Exon 3
gtaaatgccagcccctgcctcaagtaacaattgattctttttgtgtgctctcccaggtctaccctgaactgcaga base pairs
catttacggtcggggacggagttcattgttaactaagaaaaacacacgagagggtccagatgggacttgacgtct 301 to 375
                                          sgRNA, p05
tcaccaatgtggtagaagccaaccaaccagtgaccatccagaactggtgcaagcggggccgcaagcagtgcaaga base pairs
agtggttacaccatcttcggttggttggtcactggtaggtcttgaccacgttcgccccggcgttcgtcacgttct 376 to 450

cccatccccactttgtgattccctaccgctgcttaggtgagccggccggccgtggggctggtgttgattggggc base pairs
gggtaggggtgaaacactaagggatggcgacgaatccactcggccggccggcaccccgaccacaactaaccccg 451 to 525

ctggtcttgagggaagaaaaagaggatgctcctgttaggtcacatacacagacttgttcttcagcacattgccac base pairs
gaccagaactcccttctttttctcctacgaggacaatccagtgtatgtgtctgaacaagaagtcgtgtaacggtg 526 to 600

tctgtgttgtactgtgttttggactcttgcagttacattctgtgcactgaccctataggagcagtattttgagt base pairs
agacacaacatgacacaaaacctgagaacgtcaatgtaagacacgtgactgggatatcctcgtcataaaaactca 601 to 675
                                                     p06
tccctgcctcagaatgaatttacccagggtgtatattgaaattacaaattcctgggccagttccaggactcctga base pairs
agggacggagtcttacttaaatgggtcccacatataactttaatgtttaaggacccggtcaaggtcctgaggact 676 to 750

atgaaaaatgcctatagtagcggatccgggaattcttattttaccgtatcgcatagatgattctcatgaacaggg base pairs
tactttttacggatatcatcgcctaggcccttaagaataaaatggcatagcgtatctactaagagtacttgtccc 751 to 825
                                                  p04
```

**Figure S1.** sgRNA and PCR primers in reference to APP cDNA.

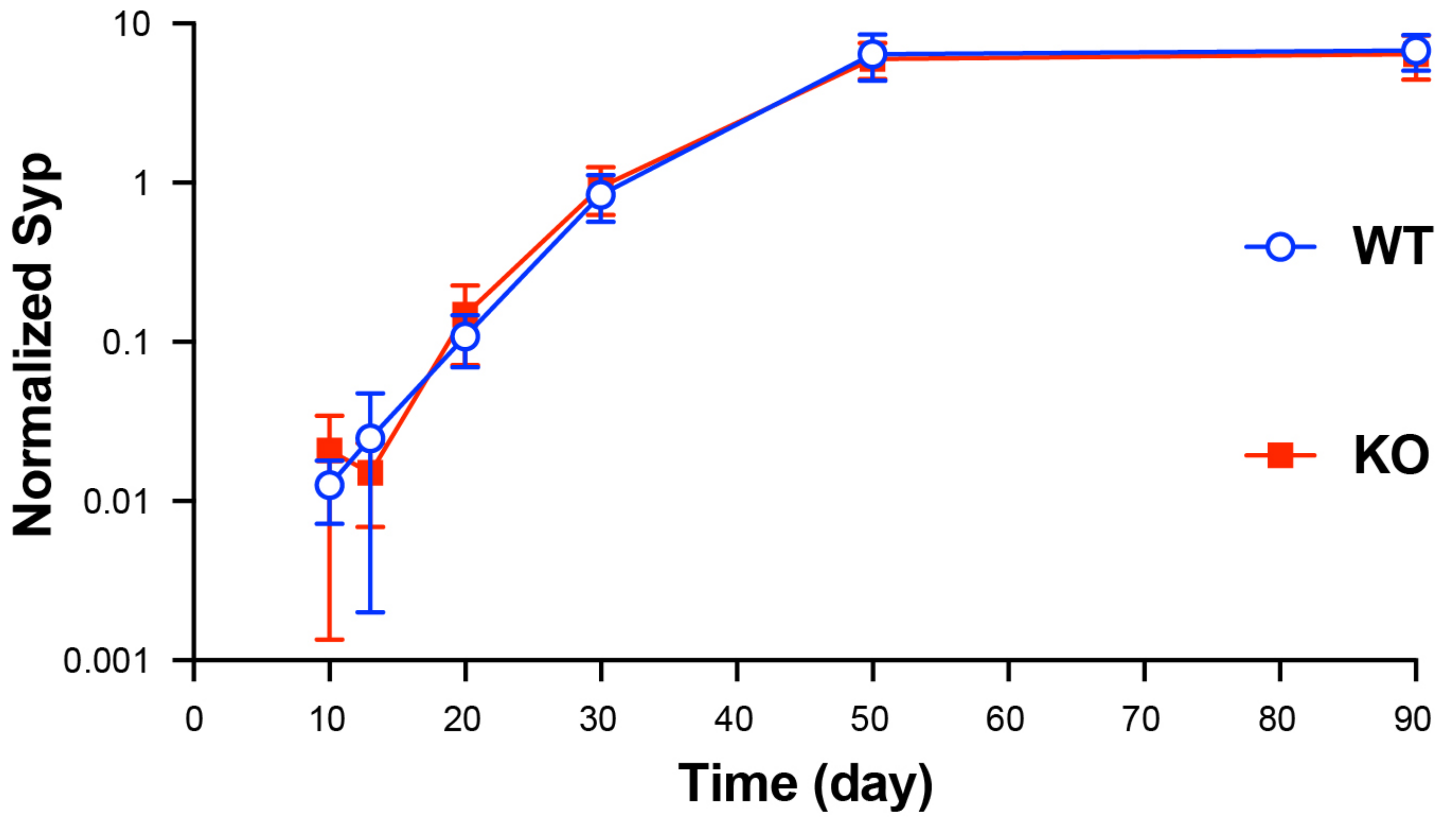


**Figure S2.** RT-qPCR detection of synaptophysin expression in hiNs. Mean and error plot of Syp by normalizing C(t) values (by β-actin, CypA, and Ppp1ca) shows no significant difference of Syp expression between WT (blue cycle) and KO (red square). All n = 9 samples (i.e., 3 samples per batch and 3 batches per group) for WT and KO at every dates. Two-tailed Student's *t*-test for pair-wise comparison at all dates, all $p > 0.05$.
